# A Gentle Introduction to Spatial Transcriptomic Analysis with 10X Visium Data

**DOI:** 10.1101/2025.05.01.651786

**Authors:** Jessica Gillespie, Juan Xie, Kyeong Joo Jung, Gary Hardiman, Maciej Pietrzak, Dongjun Chung

## Abstract

Spatial transcriptomics (ST) combines single-cell RNA-seq gene expression data with spatial coordinates to provide an accurate, 2D picture of gene expression across a tissue sample. With this technology, we can discover detailed RNA localization, study development, investigate the tumor microenvironment, and create a tissue atlas. Full ST analysis requires several steps, however, as this protocol aims to be a simple introduction to the analysis process, only the foundational steps of clustering, spatially variable gene discovery, and cell-cell communication are presented, focusing on data obtained from the 10X Genomics Visium platform. An expanded protocol with full code is available at https://github.com/j-gillespie-dna/STanalysis.git

## 1. Introduction

As biological technologies continue to advance, analysis methods that bring novel insight to the field are rapidly developing, including new ways to use already familiar techniques. RNA sequencing (RNA-seq) began with bulk methods that averaged expression across a sample. The next iteration used dissociation to disrupt tissue into single cells that could be sequenced individually (single-cell RNA-seq; scRNA-seq) to inform which genes were expressed in tandem in the same cell. Most recently, spatial information has been added to the analysis so now gene expression across a tissue sample can be reconstructed to give a 2D (or even 3D) picture. This new field is called spatial transcriptomics (ST)^1^.

ST analysis has a wide variety of applications. Starting at the sub-cellular level, imaging methods are able to localize RNA to specific parts of a cell to illuminate the structured handling of gene expression. At the cellular level, the distance between cells and the expression of ligand-receptor pairs are used to predict possible cell communication with greater accuracy. Finally, tissue-level datasets can inform the cellular makeup of a tumor sample or allow us to create a highly detailed tissue atlas with incredible resolution^2,3^.

There are several methods for capturing RNA expression and spatial information from a tissue sample, one of the more popular options being 10X Genomics’ Visium platform. The Visium platform allows for capture of close to single-cell resolution whole transcriptome RNA-seq data while also noting the spatial position of cells in a tissue. Briefly, fresh frozen or formalin-fixed paraffin-embedded (FFPE) samples are fixed to a slide and stained with immunofluorescent (IF) markers or Hematoxylin and Eosin (H&E) stain before the slide is imaged. Tissue on the slide is then hybridized with whole-transcriptome probes and the slide is placed into a 10X Genomics CytAssist machine where areas of the tissue are captured and sequenced. Genomics’ Space Ranger software then converts raw data into an expression matrix and spatial coordinates. This platform is currently a popular choice and therefore this protocol will focus on analyzing data collected this way.

Although ST analysis pipelines may involve numerous steps, this chapter highlights three core components that are consistently implemented in widely used workflows.. The first is clustering which uses spatial and RNA-seq data to group sequencing spots by common gene expression profiles. These clusters can then inform tissue architecture and be used for spatial domain prediction. Next, spatially variable gene expression detection highlights which genes may vary in expression by location revealing structural domains or tissue heterogeneity^4^. Cell-cell communication allows for the prediction of communication networks across a sample by assessing the distance between expression of ligand-receptor pairs in cells.

This protocol will focus on ST analysis with the three steps mentioned above. The software used at each step is one of a myriad of options and was chosen for ideal dataflow and ease of use. All are available as R packages therefore a familiarity with basic R functionality such as installing and loading R packages, handling R datatypes, and loading and saving data in R is advised.

## 2. Materials

### 2.1 Computer Infrastructure

As R will be the main working environment any compatible R operating system is sufficient. ST datasets can be quite large (10’s of GB) and saving data at several stages in the process may be desired. A multi-core processor for faster analysis may be beneficial at some steps, though not required. Finally, a fairly large amount of RAM may be required (>32GB). High Performance Computing services are available allowing users to upload their data to a cluster.

### 2.2 Software

R is the coding environment in which the analysis will take place. R can be run via the command-line or used in the RStudio GUI. Packages can then be installed from within R. Version numbers used in this protocol are provided but these specific versions are not mandatory.

R (v4.4.2) https://www.r-project.org/

RStudio (RStudio 2024.12.0+467 “Kousa Dogwood”) https://posit.co/downloads/

Seurat^5^ (v5.2.0) https://satijalab.org/seurat/

SPARK-X^6^ https://xzhoulab.github.io/SPARK/

CellChat^7^ (v2.1.0) https://github.com/jinworks/CellChat

## 3. Methods

### 3.1 Collect necessary files

#### 3.1.1 Space Ranger output

After sequencing is completed, 10X Visium data will be provided as either base call files (.BCL) or FASTQ files, either of which can be used for input to 10X Genomics’ Space Ranger software. The output of Space Ranger is the input for our first analysis method so we will start there. Space Ranger provides a variety of files containing data and various metrics on the samples and sequencing run. We are interested in one file and one folder. We need the filtered feature-barcode matrices: hdf5 which holds the barcodes, features, and RNA count data. We will also need the spatial folder which contains data pertaining to the image that accompanies the sample. For a full picture of output file structure, see the 10X Genomics website (https://www.10xgenomics.com/support/software/space-ranger/latest/analysis/outputs/output-overview).

#### 3.1.2 Reference data

In order to mark cell types, we need to compare gene expression to that of already determined cells. This is frequently done with a reference dataset. While the software mentioned here has some reference sets built in, there are others available for download. These can be used after the clustering step for cell-type annotation and in deconvolution. Some annotated datasets are available from the HuBMAP Consortium (https://azimuth.hubmapconsortium.org/).

#### 3.1.3 Example Datasets

10X Genomics has freely available many datasets produced with their technology for users to download and analyze. These are useful for practice and learning. A mouse anterior brain dataset will be used in this chapter and is available here (https://www.10xgenomics.com/datasets/mouse-brain-serial-section-2-sagittal-anterior-1-standard).

### 3.2 General Workflow

Figure 1. shows the general workflow of this pipeline along with the software for each step. The workflow is presented linearly here although the steps after clustering may be done in any order and several are optional depending on the desired analysis.

**Figure 1.**
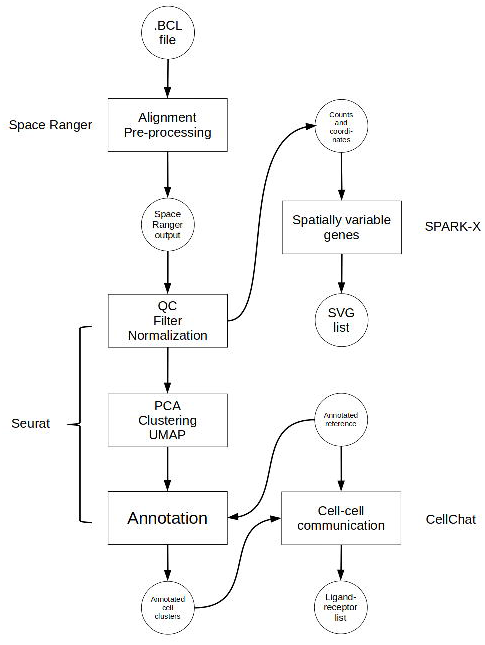
Flowchart illustrating a spatial transcriptomics analysis protocol

### 3.3 QC, Clustering, and Annotating with Seurat

First, we will import the necessary Space Ranger output and perform some QC on the gene expression data. Then the gene expression and spatial data will be used to group (cluster) cells. Finally, we will annotate the clusters by cell type.

#### 3.3.1 Importing and QC

A Seurat object must first be constructed from the hdf5 file and spatial folder. This object is a special data type, holding all of the information about the sample and any additional information as a result of the analyses performed here. Image data can be read from the spatial folder via the “Read10X_Image” command and stored as a variable. This variable can then be used in the “Load10X_Spatial” command along with the hdf5 filename to load the data.

A few plots may be viewed for quick data exploration. The “VlnPlot” and “SpatialFeaturePlot” functions with the “nCount_Spatial” option display the distribution of spatial features in the sample. This demonstrates that molecular counts vary across the sample as a result of technical variations but also tissue type. We may want to remove feature counts that are exceptionally low or extremely high. Using the “PercentageFeaturesSet” command with a pattern matching “^mt-” (often the designation for mitochondrial RNA) and a “VlnPlot” selected for this feature allows us to visualize the percentage mitochondrial RNA counts in samples. It is suggested to eliminate cells with high amounts of mitochondrial RNA as this will contribute noise to expression data. There is no definitive rule or cutoffs for this step and largely depends on the researcher’s evaluation of the data. Based on **Figure 2**, we use the “subset” function to select samples with a mitochondrial count < 30 and a feature count < 50,000.

**Figure 2.**
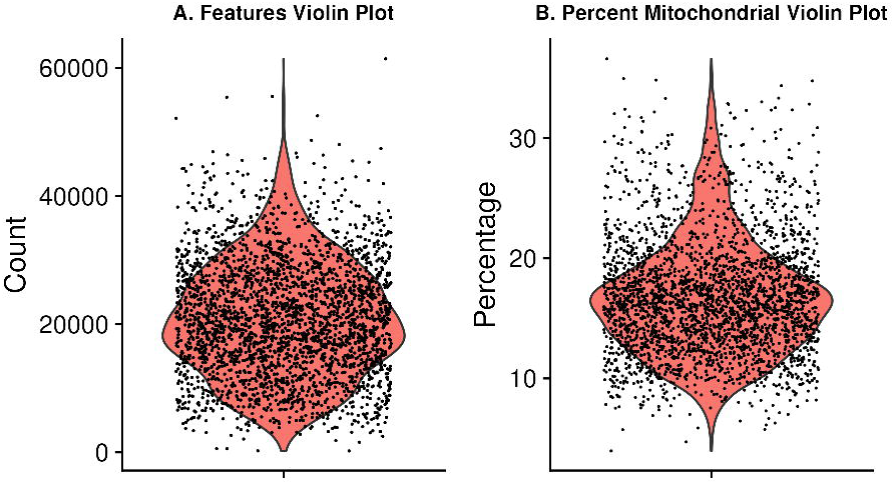
Violin plots showing spatial data QC metrics of A) Feature Counts and B) Percentage of mitochondrial RNA present

Next, we need to normalize the data using “SCTransform”. Normalization is a common step to remove from samples technical variation that may interfere with downstream analysis. Many normalization methods exist, most often log normalization is used with RNA-seq. Log normalization assumes all cells have the same number of RNA molecules, which is often far from the case and would therefore require more steps to account for this assumption. Seurat has its own method called “SCTransform” which combines the normalization and correction steps into a single command.

#### 3.3.2 Visualizing Gene Expression

Now that we have normalized gene expression, we can visualize this on the tissue sample image by again using the “SpatialFeaturePlot” only this time, providing the names of a few genes of interest. Depending on the tissue under study and our goals, it may be interesting to pick a marker that can distinguish the anatomical structure of the sample. This plot overlays the expression of chosen genes over the provided tissue H&E image. In **Figure 3**, we can see that the expression is heterogeneous for the genes shown. In the next section, we will use a method to find out which expressions vary significantly across the tissue. Note, these images are ggplot images and can therefore be customized using ggplot options.

**Figure 3.**
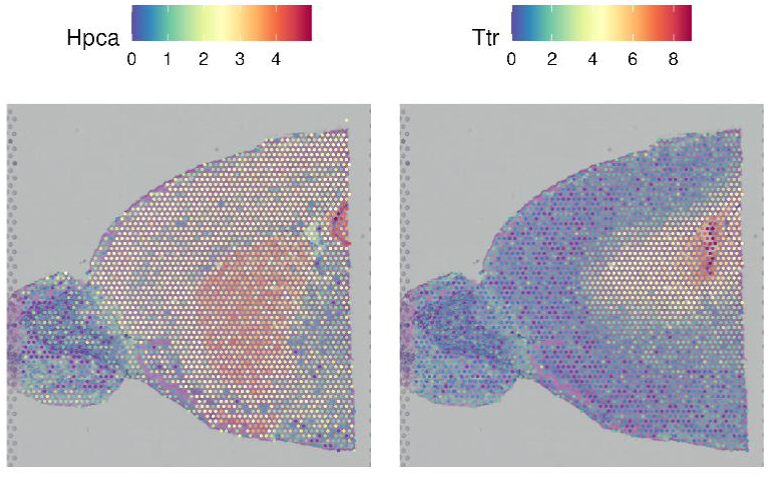
Normalized gene expression overlayed on a spatial H&E image.

#### 3.3.3 Dimensional Reduction and Clustering

When clustering, any number of variables can be considered. However, it is not necessary to include ALL variables as some contribute very little overall or even add noise. Dimension reduction pairs down the parameter list to only those that contribute meaningfully to the analysis. This is achieved here by using “RunPCA” on the transformed data. Now we can group cells using “FindNeighbors”, “FindClusters”, and “RunUMAP”. Briefly, Seurat uses a graph-based approach to clustering starting with a K-Nearest Neighbors graph based on the PCA space, then uses a Louvain algorithm to cluster the cells before a final non-linear dimensional reduction is performed with UMAP.

#### 3.3.4 Visualizing Clusters

There are a few options for visualization of our clusters. First, a traditional UMAP plot showing our clustering results with this “DimPlot” function (**Figure 4a**). “SpatialDimPlot” will show the distribution of identified clusters by overlaying them onto our tissue image. Giving “SpatialDimPlot” a few cluster identities allows us to highlight the location of selected clusters (**Figure 4b**).

**Figure 4.**
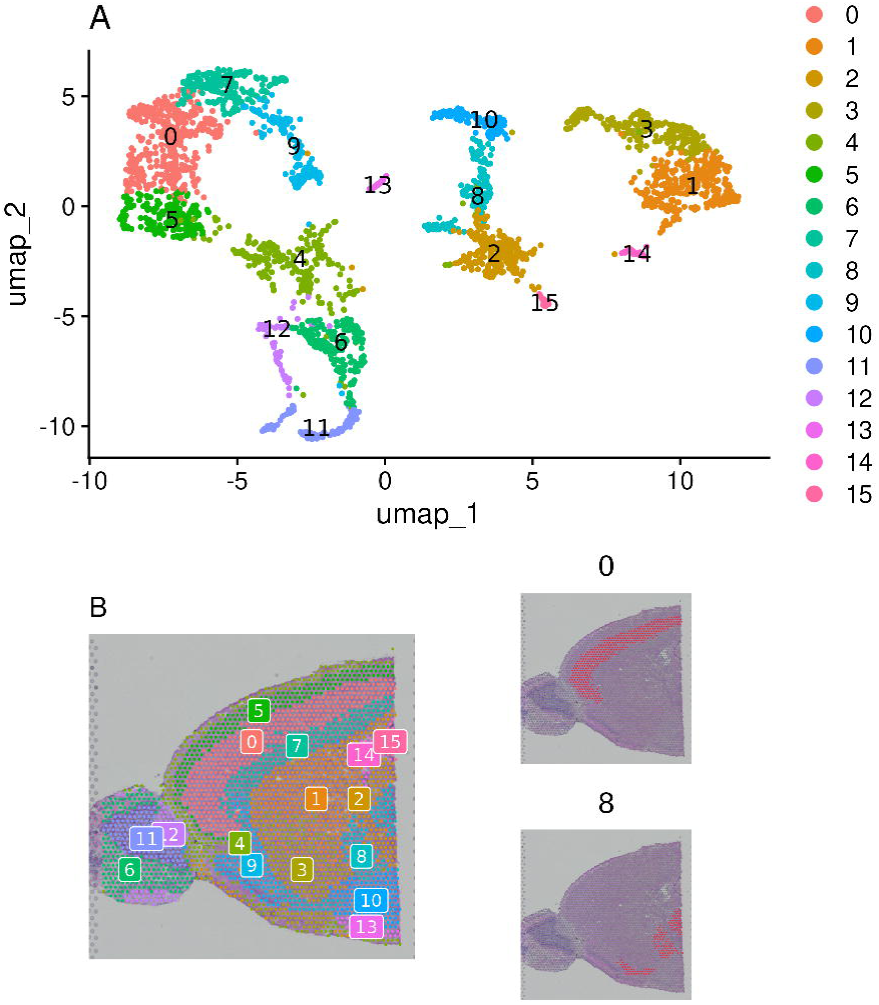
Spatial clustering results shown in A) UMAP form and B) overlayed on the spatial H&E image for all clusters (left) and highlighting chosen clusters (right)

#### 3.3.5 Spot Annotating

A final step required is to annotate our clusters by assigning them cell types. This process is not single-cell resolution as our clusters consist of “spots” that contain several cells each (a characteristic of the 10X Visium platform used here). Deconvolution is needed to estimate individual cell types in each spot and is a separate step. While there are automated methods to do this, they are far from perfect, and the best way to annotate your samples is to have domain knowledge of markers that indicate cell types. If particular markers are known, expression can be viewed by cluster via a violin plot to assist in confirming. Several methods exist to perform this step. SingleR^8^ with celldex^8^ is a popular method. Another method is Azimuth^9^. Here, we will use Sc-Type^10^. The “run_sctype” function has built-in references if we provide “Brain” as the tissue type. Note that this reference is currently only for human tissue so the match will not be perfect. We can then show the new cluster assignments using “DimPlot” and “SpatialDimPlot” (**Figure 5**). Importantly, while sc_type can annotate cells for us, this should not be used as the definitive cell type. Automated cell typing is, at this time, not exact. Correct annotation requires domain knowledge of cell type markers for the tissue in question and should be carefully considered.

**Figure 5.**
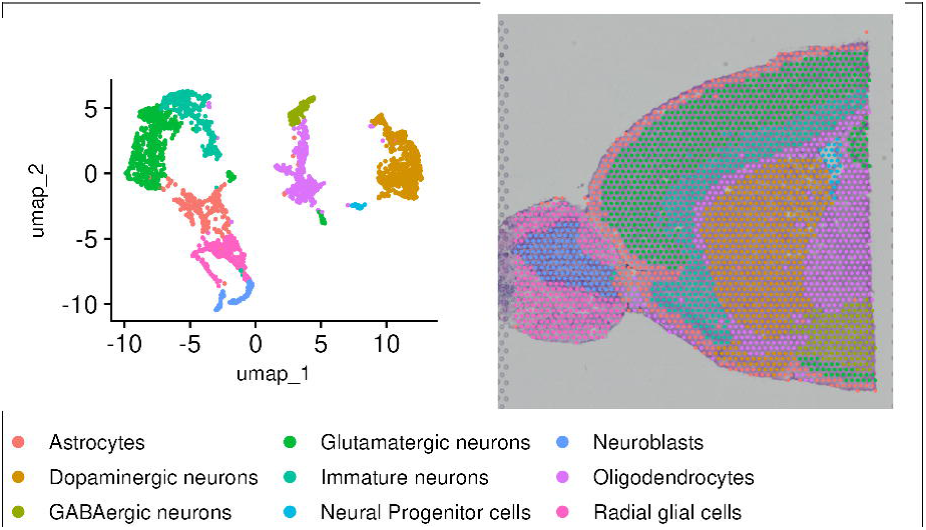
Annotated clusters shown in UMAP form (left) and overlayed on the spatial H&E image (right)

#### 3.3.6 Saving the Seurat Object

Finally, we save the Seurat object with “saveRDS”. This file can be loaded again to add other annotations or create more figures without having to repeat the analysis. This same object will also serve as the starting point in subsequent steps.

### 3.4 Spatially Variable Genes (SVGs) Discovery with SPARK-X

Now that we know our gene expression across the sample, are there genes whose expression varies across the sample in a significant way? This information can be used to study tissue structure, assess spatially-related gene expression, and determine cell-cell communication (although we will look at this specific topic in a later section). There are again many software options for this step using a variety of methods. Some use a Gaussian process (SpatialDE^11^, nnSVG^12^, SOMDE^13^) that provides good results but scales significantly with dataset size. SPARK-X uses non-parametric methods to evaluate the independence of expression of a gene with location on the tissue. This approach requires significantly less RAM, even for large datasets. When finished with this step, we will have data on which genes vary in their expression across the tissue sample and a figure to show as much.

#### 3.4.1 Importing and Preparing Data

Using “readRDS” we can load in our saved Seurat object. Using “GetAssayData” and “GetTissueCoordinates” from Seurat, we can create two variables that contain the raw expression counts and the spatial coordinates respectively. Note if you have not yet removed mitochondrial genes, this should be performed before running this analysis (see Section 3.3.1).

#### 3.4.2 Running SPARK-X

With a single “sparkx” command, spatial information and gene expression are used to determine which gene’s expression differs by tissue area.

#### 3.4.3 Visualizing SVGs

Now that we have a list of spatially variable genes, we can look at them across our tissue sample using the Seurat “SpatialFeaturePlot”. Here we look at the top 10 most significantly spatially variable genes (**Figure 6**).

**Figure 6.**
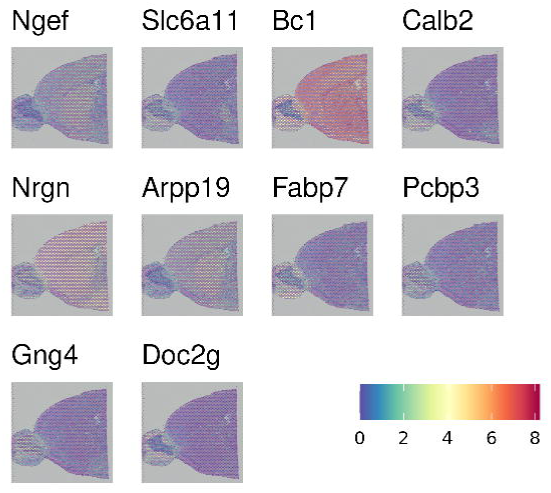
Gene expression of the 10 most significantly spatially variable genes

#### 3.4.4 Saving SVG Data

We can save our SPARK-X output to an rds file for quick loading to look at specific data or create new figures.

### 3.5 Cell-Cell Communication with CellChat

Cells communicate by releasing and receiving molecular messages called ligands. Ligands bind to receptors, often found on cell surfaces. This binding then activates a reaction or pathway that leads to a cell function. Due to decades of research, we know which ligands typically bind to which receptors. Thanks to the RNA-seq data for our sample, we know which cells are expressing either the ligand or receptor in a canonical communication pair. As we also have spatial data available, we can use statistics to predict which cells are most likely communicating with one another based on ligand-receptor gene expression and physical distance from one another.

One factor that heavily influences the results of any CCC analysis is the ligand-receptor database used^14^. Indeed the same method can return two drastically different results merely by working from two different ligand-receptor pair lists. CellChat has its own database, CellChatDB, which is manually curated and takes into account things like subunit structure and regulatory pathways. The database is not exhaustive, however, and some ligand-receptor pairs may be missing. Users are able to update the database following the instructions here https://htmlpreview.github.io/?https://github.com/jinworks/CellChat/blob/master/tutorial/Update-CellChatDB.html

Another important factor is your annotations previously added in the Clustering step. CellChat will group samples by these annotations to determine which groups are potentially communicating with one another. Be sure your sample is well annotated to obtain the most accurate communication network.

#### 3.5.1 Importing Data and Setting Compute Options

Once again, we can load in the rds of the Seurat object from clustering. No SVG data is saved in the object from the previous section but it is not needed here. This step is very time intensive, potentially requiring several hours on a personal or office machine. Two options can speed up the process, however.

R generally reserves only a small amount of RAM for calculations (~500MB) which is easily not enough for this step. We can set an option for the future.globals.maxSize to take advantage of a larger amount of RAM. The exact number will be system-dependent, here we used 32GB of RAM by providing the value 32000*1024*8. We can take advantage of a multicore processor by utilizing the future::plan function to set the number of cores we would like to use for CCC.

#### 3.5.2 Preparing CellChat Object and Database

There is a single command needed to create a CellChat object from a Seurat object. Preparing the database is somewhat more complicated. CellChat has two main databases, one for human and one for mouse. The databases contain a variety of communication modes including secretionary, cell-cell contact, and non-protein signaling. Depending on our cell types and desired research direction, these databases can be subset to focus only on specific interaction lists which may decrease processing time and simplify results. Finally, we need to convert pixels in the H&E image to real-world measurements using the “scalefactors_json.json” file provided in the Space Ranger output and the spot size of the 10X Visium assay (usually 55μm). A CellChat object can then be created with the aptly named “createCellChat” function.

#### 3.5.3 Running CellChat

Before finding all ligand-receptor pairs, over-expressed genes and interactions are found to infer the cell state-specific communications using “identifyOverExpressedGenes” and “identifyOverExpressedInteractions” respectively. If the sequencing depth is shallow, we can use “smoothData” to smooth genes’ expressions based on neighbors’ validated and high-confidence protein-protein network. If smoothing is used, the parameter “raw = FALSE” should be added to the “computeCommunProb” function.

CellChat offers two options for calculating ligand-receptor pairs, “triMean” which produces fewer but stronger interactions, and “truncatedMean” to identify weaker signaling. We use our chosen option in “computeCommunProb” to run the calculation. This function can take several hours to run, especially if using a personal computer instead of cluster. Thankfully, a progress bar is conveniently displayed.

After our list is obtained, we filter out groups that have only a few cells on which to predict communication with “filterCommunication”. Next, “computeCommunProbPathway” will use the ligand-receptor interactions to calculate the communication probability on a signaling pathway and can then aggregate an entire communication network using “aggregateNet”. As the previous step takes a significant amount of time, it is best to save our analysis as an rds file.

#### 3.5.4 Visualizing CCC

There are many figures and graphs that can be created for CCC analysis. First, using “netVisual-circle”, we can view communications between each cell group of cells from the clustering step. The vertex and line weights can be set to a number of parameters, including the number of interactions and the strength of the interactions (**Figure 7a and b**). The same information can alternatively be viewed as heatmaps using “netVisual_heatmap” (**Figure 7c and d**).

**Figure 7.**
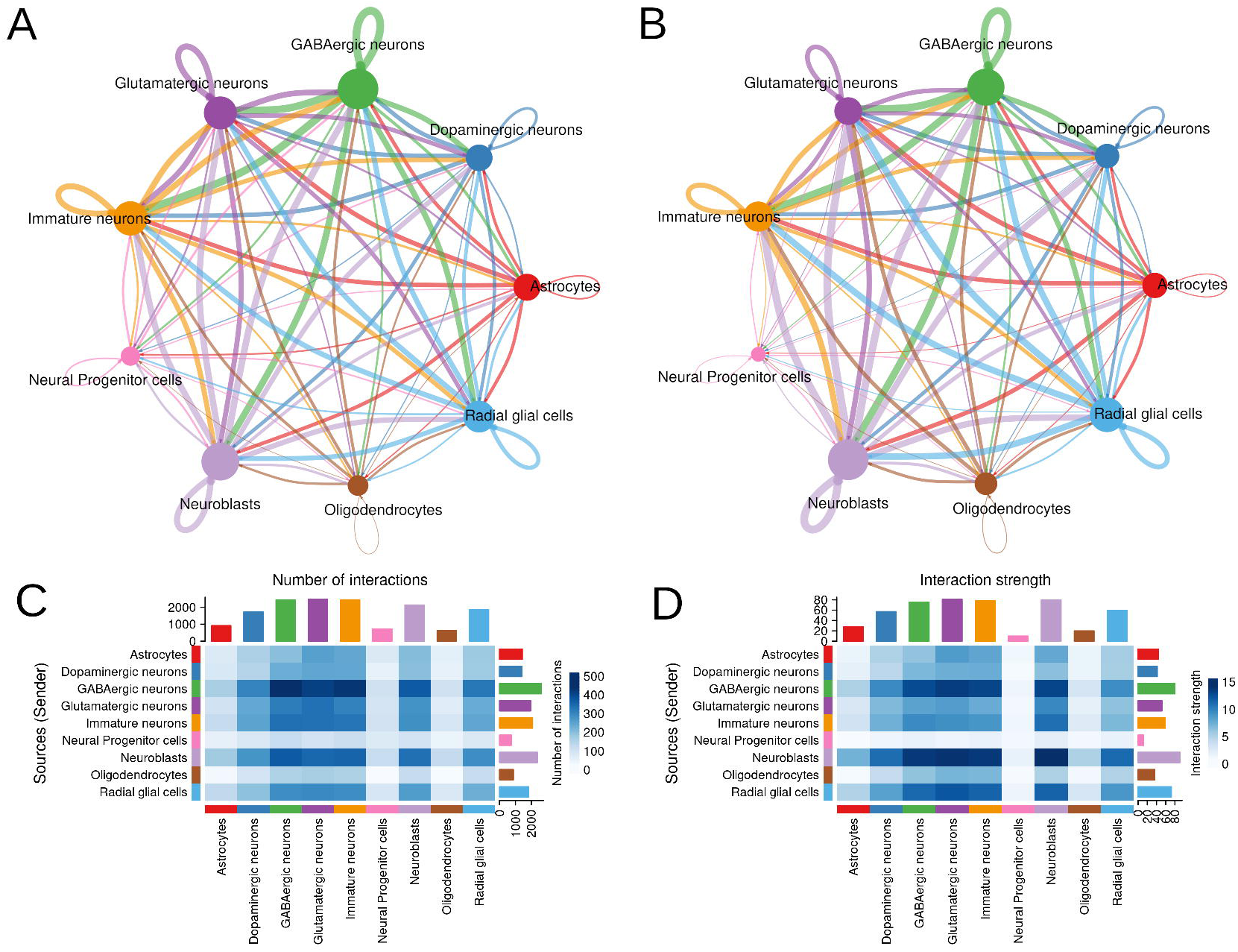
Cell-cell communication network by clustered cell types. Emphasized by A) number of counts and B) strength of interaction in network form and in heatmap form (C and D).

If we are interested in a particular signaling pathway, a list of communication probabilities is available in the CellChat object metadata. We can look at the probability of interactions between cell groups involved in the NOTCH pathway with the “netVisual_aggregate” function (**Figure 8**).

**Figure 8.**
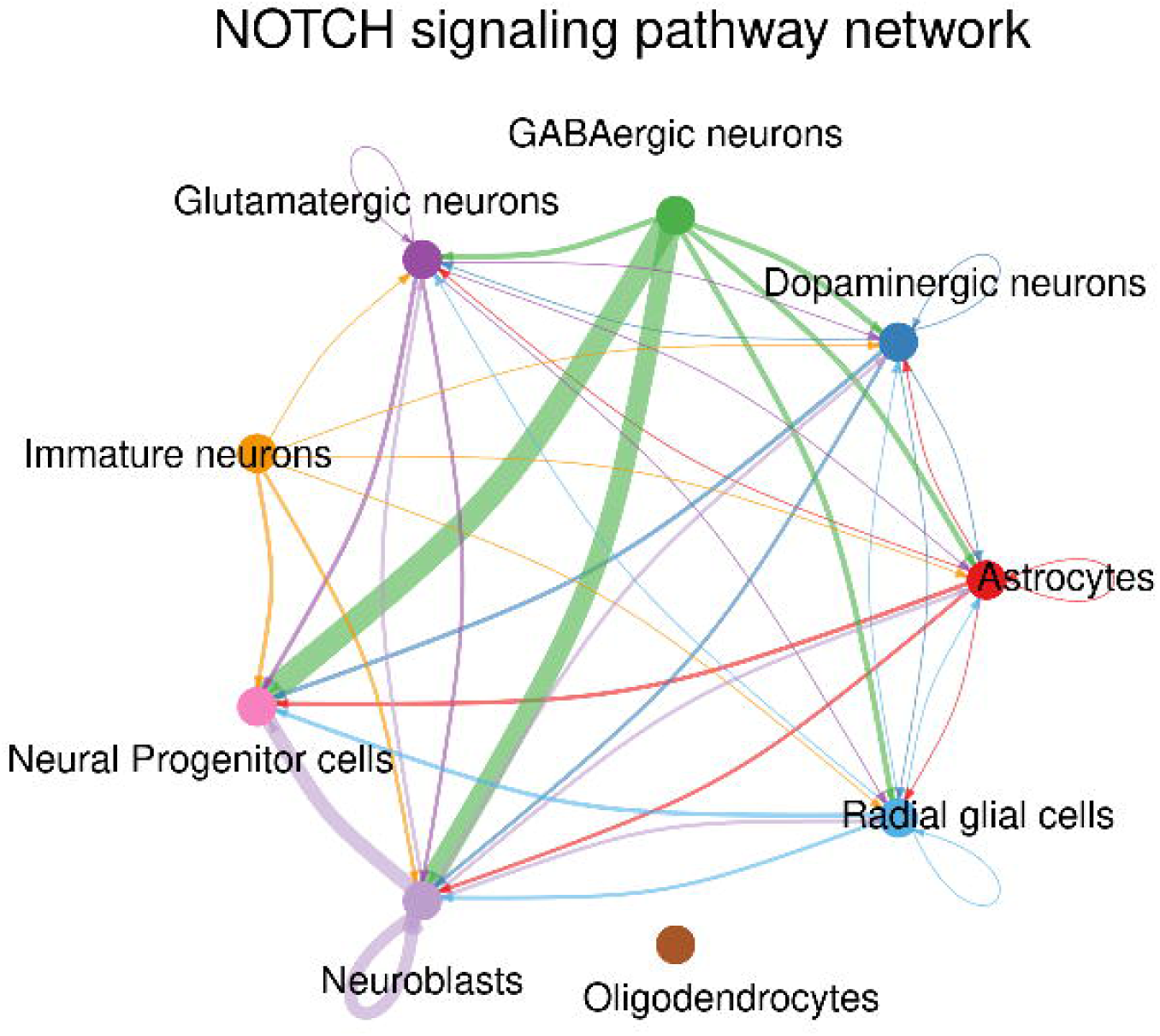
Aggregate communication probabilities between clustered cell groups.

Finally, we can explore the “centrality” of a network. This tells us which nodes are active in the most pathways and therefore could be more “important”. The calculation is done with “netAnalysis_computeCentrality” and visualized with “netAnalysis_signalingRole_network” (**Figure 9**).

**Figure 9.**
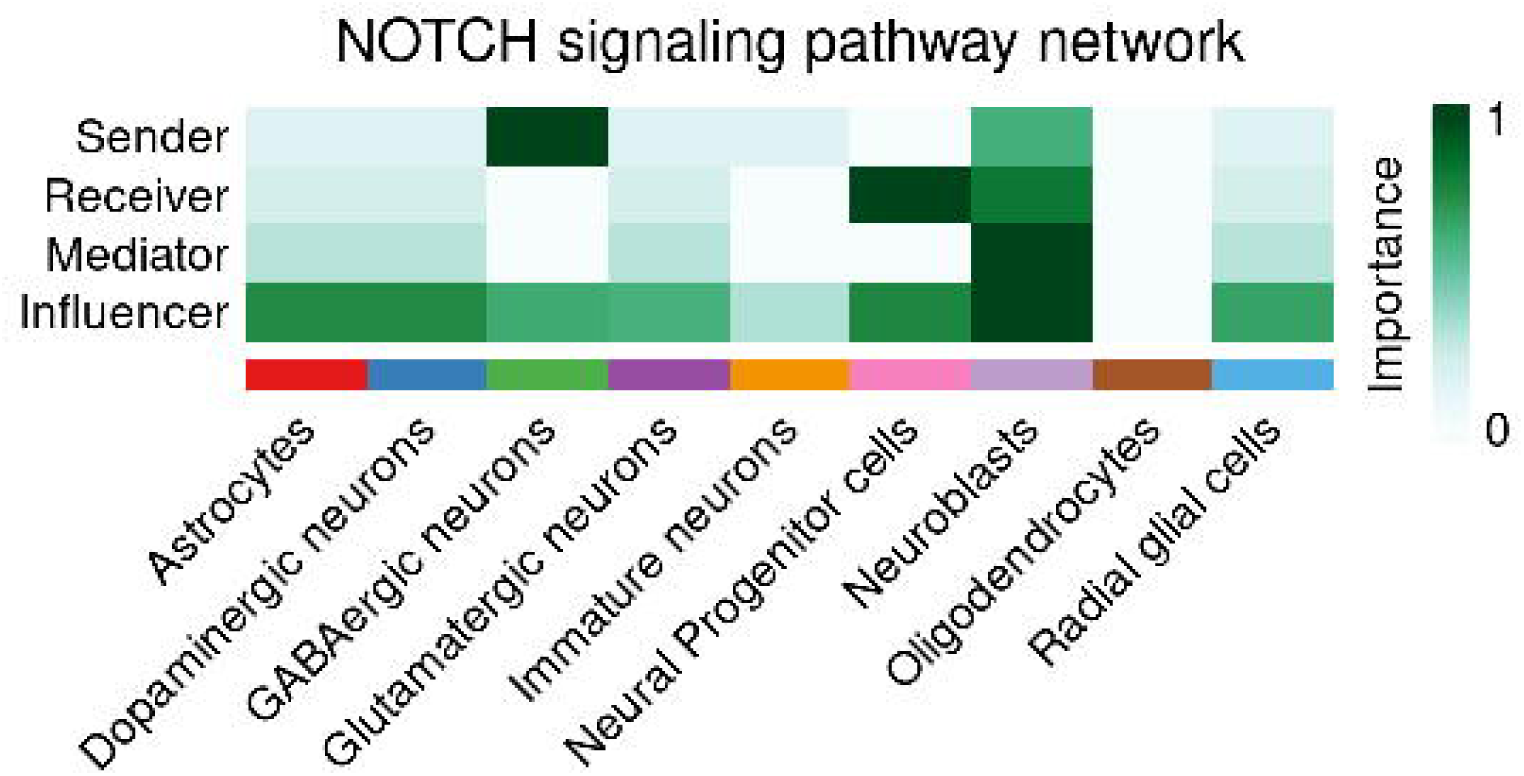
Heatmap of cell type centrality in communication network.

As we saved the CellChat object after the analysis and have not added to it since then, we do not need to save it again here.

### 3.6 Further Analysis

As this step is optional, it will not be covered here in detail. However, we did want to mention two R packages, RCTD^15^ and CARD^16^ that are useful for deconvolution. RCTD uses expression profiles from a reference dataset and supervised learning to determine cell type proportions in a spot. CARD uses a gene expression reference and spatial correlation to determine cell type at each spot across a tissue. Either package requires our Seurat object with spatial data and an annotated reference RNA-seq dataset appropriate for the tissue under study.

As we have RNA-seq data, analyses typically performed with these data can also be done here. Differential gene expression (DEG) is often performed. We can examine DEG per cell group defined in the clustering step or we can also compare expression in one cluster versus the rest of the cell population in the tissue. Seurat has methods to perform this analysis. Gene set enrichment analysis (GSEA)^17^ or a pathway analysis are also possible areas for further study.

## 4. Conclusion

Spatial transcriptomics adds another level to RNA analysis, combining gene expression and location information. While we present a simple workflow with mostly default options, there are many nuances to ST analysis. For example, many software has the option to pre-process data before clustering yet there has been evidence that pre-processing can greatly affect analysis outcome^18^. Similarly, normalization can affect deconvolution and it is suggested only raw spatial data should be used^19,20^. For SVGs and CCC, the list of results returned and their associated statistics greatly depends on the software and database choice respectively^14,21,22^. It has been demonstrated in multiple studies that analysis performance is highly dependent on the dataset/software pairing and further algorithm refinement is necessary^18–20,22,23^.

Here we focus on one platform and a handful of software for analysis but there are many options available. Visium is only one sequencing platform with other popular options being Slide-seq, MERFISH, seqFISH, Visium HD, and the emerging 10X Genomics Xenium. Each platform has its own pros and cons and offers slightly different analysis options. There are also many software options for each analysis step mentioned here. Again, each has distinct advantages and strengths depending on the dataset and sequencing platform. In the future, we would like to expand this protocol to make it applicable to a wider variety of platforms and analysis options.

